# *Ex Vivo* Culture Models *of* Hidradenitis Suppurativa for defining molecular pathogenesis and treatment efficacy of novel drugs

**DOI:** 10.1101/2021.10.11.464007

**Authors:** Kayla Goliwas, Mahendra P. Kashyap, Jasim Khan, Rajesh Sinha, Zhiping Wang, Allen S W Oak, Lin Jin, Venkatram Atigadda, Madison B. Lee, Craig A. Elmets, M Shahid Mukhtar, Chander Raman, Jessy S. Deshane, Mohammad Athar

## Abstract

Hidradenitis suppurativa (HS) is a complex inflammatory and debilitating skin disease for which no effective treatment is available. This is partly because of the unavailability of suitable human or animal models with which exact pathobiology of the disease can be defined. Here, we describe the development of air-liquid (A-L) interface, liquid-liquid/liquid-submersion (L-S) and bioreactor (Bio) *ex vivo* skin culture models. All three *ex vivo* platforms were effective for culturing skin samples up to day-14, with the tissue architecture and integrity remaining intact for at least 3 days for healthy skin while for 14 days for HS skin. Up to day-3, no significant differences were observed in % early apoptotic cells among all three platforms. However, an increase was observed in late apoptotic/necrotic cells in HS skin at day-3 in A-L and Bio culture of HS skin. These cultures efficiently support the growth of various cells populations, including keratinocytes and immune cells. Profiling of the inflammatory genes using HS skin from these *ex vivo* cultures showed dynamic expression changes at day-3 and day-14. All of these cultures are necessary to represent the inflammatory gene status of HS skin at day-0 suggesting that not all gene clusters are identically altered in each culture method. Similarly, cytokine/chemokine profiling of the supernatant from vehicle- and drug-treated *ex vivo* HS cultures again showed better prediction of drug efficacy against HS. Overall, development of these three systems collectively provide a powerful tool to uncover the pathobiology of HS progression and screen various drugs against HS.

## INTRODUCTION

Hidradenitis suppurativa (HS), is a chronic inflammatory skin condition with painful lesions, including abscesses, draining tracts, and fibrotic scars. This debilitating disease affects 1%–4% of the Western population ^1, 2^. So far, no effective treatments have been identified particularly for severe disease. The pathologic process of HS is thought to arise from occlusion and rupture of defective hair follicles and the subsequent release of contents including bacteria into the dermis ^3,4^. A complex inflammatory response by surrounding neutrophils and lymphocytes is known to accelerate the formation of abscess and cause the subsequent destruction of the pilosebaceous unit ^5,6^. Although, the presence of abscesses and malodorous drainage has implicated a role of microorganisms in HS pathogenesis, certain antimicrobial therapies have only been temporarily and/or partially effective. In addition, the risk of drug resistance is high, if employed on a long-term basis ^7^. Keratinocyte hyperproliferation is also a hallmark of HS ^8^. Unlike other skin diseases, the immunobiology of HS is poorly understood, which has also hindered the development of effective therapies. Studies defining the elevation of inflammatory cytokines, such as tumor necrosis factor alpha (TNF-α), interleukins (IL)-1β, IL17A and various others in HS patients provided possible therapeutic targets for emerging treatments ^9–12^. This is also based on the known success of biologicals against these targets in various other pro-inflammatory diseases ^13^. The TNF-α antagonist adalimumab is the first and only FDA– approved therapy for this disease in the USA ^13^. Lack of optimal pre-clinical models of HS has hampered the pace of development of novel therapeutics to target effectively immune dysregulation and associated pathobiology of HS disease progression ^14^.

In this study, we report the establishment of three different *ex vivo* HS skin culture models that enable efficient characterization of immune infiltrates and hyperproliferative keratinocytes in HS tissues alone and following treatment with various drugs. These models include air-liquid interface (A-L) culture, liquid submersion (L-S) culture, and culture using a three-dimensional perfusion bioreactor (Bio) platform with well-defined culture condition. L-S culture is very similar to traditional cell culture, with the entire three dimensional tissue submerged in culture media. A-L culture better mimics the native tissue environment, with the apical tissue surface exposed to air and the basal surface covered by media. Both the A-L culture and the L-S culture are static culture platforms, whereas the Bio platform allows for continuous perfusion of oxygen/nutrients to mimic the human circulatory system. Thus, each of these methods have some benefits over others as well as some disadvantages. We demonstrate that the tissue architecture and viability of both healthy and HS skin tissues are maintained *ex vivo* in these models, over a period of 14 days. CD4^+^ and CD8^+^ lymphocyte populations, including the regulatory T cells and memory T cells, as well as keratinocytes and monocytes/macrophages could be detected and quantified by FACS analysis in these models. Importantly, we also assess the feasibility of these models to test the effects of drugs on the inflammatory cytokines/chemokine signatures and signaling pathways associated with immune dysfunction in HS. Utilization of such a multi *ex vivo* HS skin culture platform will facilitate the development of novel therapeutics and unravel unique mechanisms underlying the pathogenesis and therapeutic action of drugs.

## METHODS

### Human Tissue

The institutional Review Boards of the University of Alabama at Birmingham approved the protocols (N081204004) for obtaining discarded surgically removed skin tissues from healthy and HS (Hurley stage 2 or 3) subjects.

### Sample Processing and *Ex Vivo* Culture

Clinical specimen were divided into ~5 cm x 5 cm tissue pieces. Tissue pieces were then either (1) placed onto a 0.4 μm transwell filter (Merck Millipore) with a thin layer of extracellular matrix (ECM, 90% collagen type 1 (Advanced Biomatrix, USA) + 10% growth factor reduced Matrigel (Corning, USA)) components for stability or (2) placed into the central chamber of a polydimethylsiloxane (PDMS, Krayden, USA) bioreactor containing a mixture of ECM for structural support as previously described ^15^. The tissue/ECM volume within the bioreactor was penetrated with five 400 micron Teflon coated stainless steel wires to generate through-channels for tissue perfusion. Following ECM polymerization, the wires were removed and the through-channels were filled with tissue culture media (1:1 mixture of X-Vivo15 and Bronchial Epithelial Growth media (Lonza, USA) with antibiotics (MP Biomedicals, USA)). The bioreactor was then connected to a perfusion system, that contained a media reservoir, peroxide cured silicon tubing (Cole Parmer, USA), a collection reservoir and peristaltic pump (ESI, USA), and tissue culture media was perfused through the tissue volume for 3 to 14 days (37°C, 5% CO2), with media changed every 3 days. Following ECM polymerization, tissue culture media was added to the transwell filters and to the bottom of the well to generate liquid submersion (tissue fully submerged in media) and air-liquid (the media volume did not cover the top tissue layer) cultures. When applicable, treatment with lenalidomide (1.5 mg/mL, Selleckchem), CPI-0610 (1.5 mg/mL, Selleckchem), or vehicle control (dimethyl sulfoxide (DMSO)) occurred daily for 3 days culture. At the end of each experiment, a portion of each tissue was fixed separately for histologic processing and collagenase B (Roche, Switzerland) digestion for flow cytometry analysis. The supernatant culture media was collected for assaying cytokine/chemokines levels.

### Histologic Processing and Analysis

Following *ex vivo* culture, a portion of each cultured tissue was fixed with neutral buffered formalin, processed to paraffin, and histological sections were prepared, as previously described ^15^. 5 micron sections were stained with hematoxylin and eosin (H&E) to evaluate tissue morphology.

### Analysis of Cell Viability via Flow Cytometry

The PE Annexin V Apoptosis detection kit I (BD, Germany) was used to analyze cell viability following manufacturer’s instructions. Analyses were performed on FACSymphony A3 Cell Analyzer with FACSDiva software version 8.0.1 (BD, Germany). Data were analyzed with FlowJo 10.7.1 (Treestar, USA).

### Multi-parametric Flow Cytometry

<u>The following antibodies were used for multiparametric flow cytometry for T cell analysis:</u> Anti-CD3-Alexa Fluor 700 (Clone: UCHT1); anti-CD4-FITC (Clone: RPA-T4); anti-CD69-BV563 (Clone: FN50); anti-CD25-BV510 (Clone: M-A251) from BD Biosciences (Germany). Anti-CD3-PE-Cy7 (Clone: UCHT1); anti-CD8-APC (Clone: 53-6.7); anti-CD4-PE-Cy7 (Clone: SK3); anti-FoxP3-PerCP-Cy5.5 (Clone: PCH101); anti-CD8-FITC (Clonse: SK1); CD-127-APC (Clone: eBioRDR5) from eBioscience (Thermo Fisher, Germany). Anti-Ki-67-Dylight350 (Clone: 1297A) from Novus (USA). Anti-CD45-APC-Cy7 (Clone: 2D1); anti-CCR7-Pacific Blue (Clone: G043H7); anti-CD45RA-BV510 (Clone: HI100); anti-CD45RO-PerCy-Cy5.5 (Clone: UCHL1); anti-CD62L-BV650 (Clone:DREG-56); anti-CD103-PE (Clone:Ber-ACT8); anti-CD154-PE/Dazzle (Clone:24-31) from Biolegend (USA). <u>The following antibodies were used for multiparametric flow cytometry for analysis of resident immune and structural cells:</u> Anti-CD64-PerCp-eFluor710 (Clone: 10.1); anti-CD11b-APC-Cy7 (Clone: ICRF44); anti-HLA-DR-FITC (Clone: LN3); anti-EpCAM (CD326)-Alexafluor 594 (Clone:9C4); anti-CD14-PE-Cy7 (Clone:61D3); anti-CD163-PE (Clone: GHI/61); anti-CD49f-PE (Clone: GoH3); E-Cadherin-PerCP-eFluor 710 (Clone: DECMA-1); CD16-FITC (Clone: eBioCB16) from eBioscience (Thermo Fisher, Germany). Anti-CD45-Pacific Blue (Clone: HI30); CD15-Alexa Fluor 700 (Clone: WCD3) from Biolegend (USA). Anti-PanCytokerain-APC (Clone: C-11) and anti-Ki-67-Dylight350 (Clone: 1297A) from Novus Biologicals (USA). The Foxp3 / Transcription Factor Staining Buffer Set (Thermo Fisher, Germany) and the Cytofix/Cytoperm Fixation/Permeablization kit (BD, Germany) were used according to the manufacturer’s protocol to stain for intracellular molecules (intranuclear and cytoplasmic molecules, respectively). Analyses were performed on FACSymphony A3 Cell Analyzer with FACSDiva software version 8.0.1 (BD, Germany). Data were analyzed with FlowJo 10.7.1 (Treestar, USA).

#### Expression profiling of inflammatory markers

To assess the expression of inflammation related genes, we employed TaqMan based Human Inflammation Open Array panel (Cat # 4475389, Thermo Fischer Scientific, USA). This panel contained 607 targets, 586 inflammatory genes and 21 endogenous control. For this, total RNA from tissues from various *ex vivo* cultures of HS skins was isolated using Trizol reagent (Cat#15596018, Ambion). A total of 2μg of RNA was reverse transcribed into cDNA using SuperScript® VILO™ cDNA Synthesis Kit (Cat #11754250, Life Technologies, USA). Following the reverse transcription reaction, the mixture was denatured by incubating it at 85 °C. Pre-amplification of cDNA using custom TaqMan preamp primers pool (Cat#4441856, Thermo Fischer Scientific, USA) was performed in 25 μl total volume containing 12.5 μl of TaqMan PreAmp master mixture (Cat#4391128), 2.5 μl PreAmp primer pools, 2.5 μl reverse transcription product, and the remaining volume was adjusted by nuclease-free water. Pre-amplification thermal cycling conditions include incubation of products at 95 °C for 10 min followed by 12x cycles at 95 °C for 15 s and 4 min at 60 °C. The pre-amplification products obtained after the cycling were incubated for 10 min at 99.9 °C. 20 fold diluted pre-amplification product was used in final amplification reaction as per the manufacturer’s instructions. OpenArray chip containing the pre-coated primers for 607 targets was read on 12 K Flex RT-PCR machine (Thermo Fisher Scientific, Life Technologies Corporation, Grand Island, New York). Data analysis was performed through the online available Expression suit v 1.3 software (Thermo Fisher Scientific, Life Technologies Corporation, Grand Island, New York) using global gene normalization method. Bioinformatics analysis was carried out using Ingenuity Pathway Analysis (IPA, Qiagen) application.

### Chemokine Profiling

To identify the drug response on HS skin, multiplex cytokine/chemokine analysis was performed using the conditioned medium procured from various *ex vivo* cultures. Cytokine/Chemokine/Growth Factor 45-Plex Human ProcartaPlex™ Panel 1 (Cat# EPX450-12171-901, Thermofischer, USA) was used. Briefly, the condition medium was centrifuged at 10,000 rpm for 10 min, and stored at −80°C until assayed. Total forty five target proteins containing (i) Th1/Th2 markers: GM-CSF, IFN gamma, IL-1 beta, IL-2, IL-4, IL-5, IL-6, IL-8, IL-12p70, IL-13, IL-18, TNF alpha, LIF (ii) Th9/Th17/Th22/Treg markers: IL-9, IL-10, IL-17A (CTLA-8), IL-21, IL-22, IL-23, IL-27 (iii) Inflammatory cytokines: IFNα, IL-1α, IL-1RA, IL-7, IL-15, IL-31, TNFβ (iv) Chemokines: Eotaxin (CCL11), GROα (CXCL1), IP-10 (CXCL10), MCP-1 (CCL2), MIP-1α (CCL3), MIP-1β (CCL4), RANTES (CCL5), SDF-1 alpha, and (v) growth factors: BDNF, EGF, FGF-2, HGF, NGF beta, PDGF-BB, PlGF-1, SCF, VEGF-A, VEGF-D were assessed. 50μl of condition medium was used for multiplexing on the Luminex 200 instrument (Luminex Corporation, USA) as also described earlier ^16^.

### Statistical Analysis

Statistical analysis was performed using GraphPad Prism software (La Jolla, CA, USA). Data are presented as mean ± standard error of the mean unless indicated otherwise in the figure legend. A p-value less than 0.05 was considered to be statistically significant. A two-tailed unpaired Student’s t-test was used to evaluate statistical difference between two groups. One way ANOVA with Tukey’s multiple comparison testing was utilized to evaluate statistical difference for data with more than two groups.

## RESULTS

### Tissue Integrity

Skin tissue specimen were collected from healthy individuals (without evidence of any skin disease) and patients with HS. Three culture platforms were established and tested for *ex vivo* maintenance of tissues. These culture platforms include air-liquid interface culture (A-L), liquid submersion culture (L-S) and perfusion bioreactor culture, which we have previously utilized for human lung tissue (**Figure 1A-1C**) ^17^. Histologic analysis of tissues demonstrates that all three platforms are effective for culturing these skin samples, with maintenance of tissue integrity for at least 3 days (**Figure 2A**, left). When the culture period was further extended to 14 days, the healthy skin tissues showed some signs of loss of tissue architecture, whereas HS skin specimen maintained most of the tissue structure and tissue integrity when H & E stained cross-sections were analyzed (**Figure 2A**, right).

**Figure 1:**
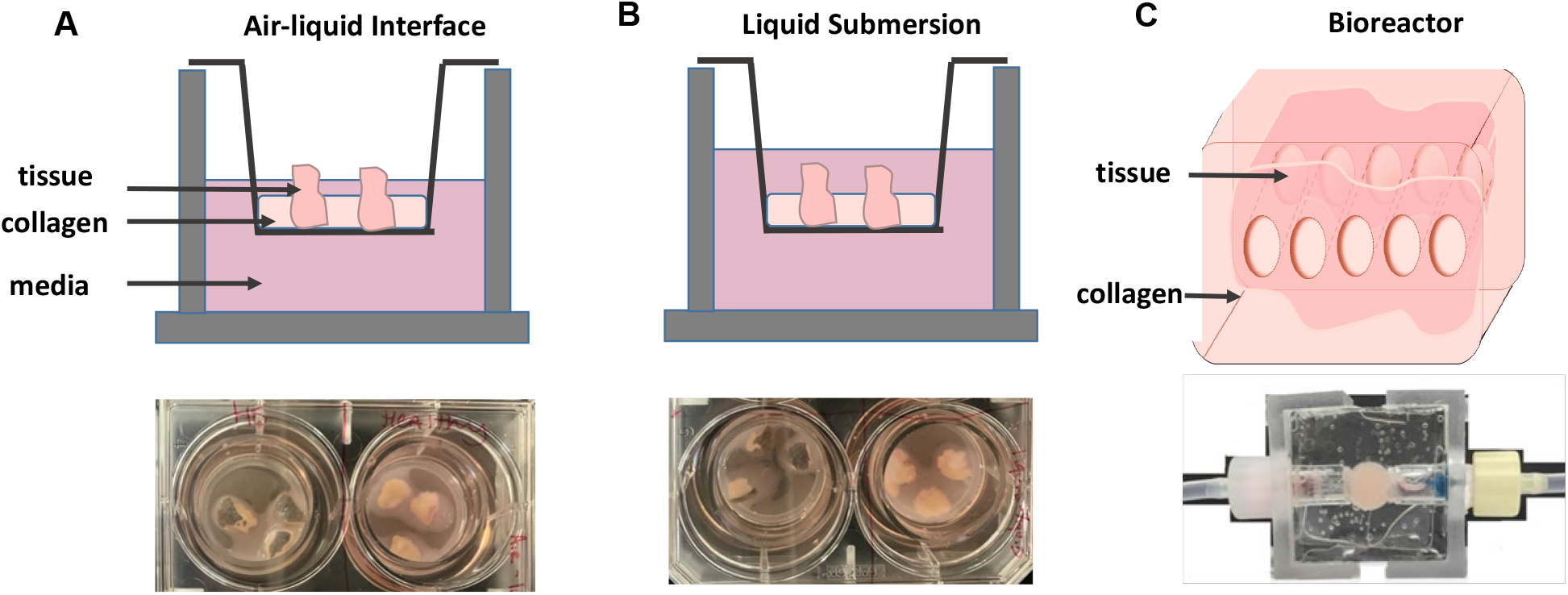
*Ex Vivo* Culture Platforms for healthy and HS skin. Schematics of three *ex vivo* culture platforms utilized to culture human skin (top) with images of each platform (bottom). **A**. Air-liquid interface (A-L) culture utilized 0.4 micron transwell filters in which an extracellular matrix (ECM) scaffold stabilized tissue specimen for static culture. Media was then added to the top and bottom chambers, but did not fully cover the tissue so that the apical surface had exposure to air. **B**. Liquid submersion culture (L-S), was set up in a similar method to A-L, with tissues stabilized within ECM on transwell filters, however media was added to cover the entire tissue. **C**. For perfusion bioreactor culture (Bio) a portion of tissue was placed into the central chamber of the polydimethylsiloxane (PDMS) bioreactor which contained an ECM scaffold for stabilization. Through-channels were generated in the tissue/ECM volume using five 400 micron Teflon coated stainless steel wires during ECM polymerization for continuous tissue perfusion. Wires were removed following ECM polymerization and the bioreactor was then connected to a perfusion system, in which a serum-free defined tissue culture media was perfused from a media reservoir through the tissue volume and collected in a collection reservoir.

**Figure 2:**
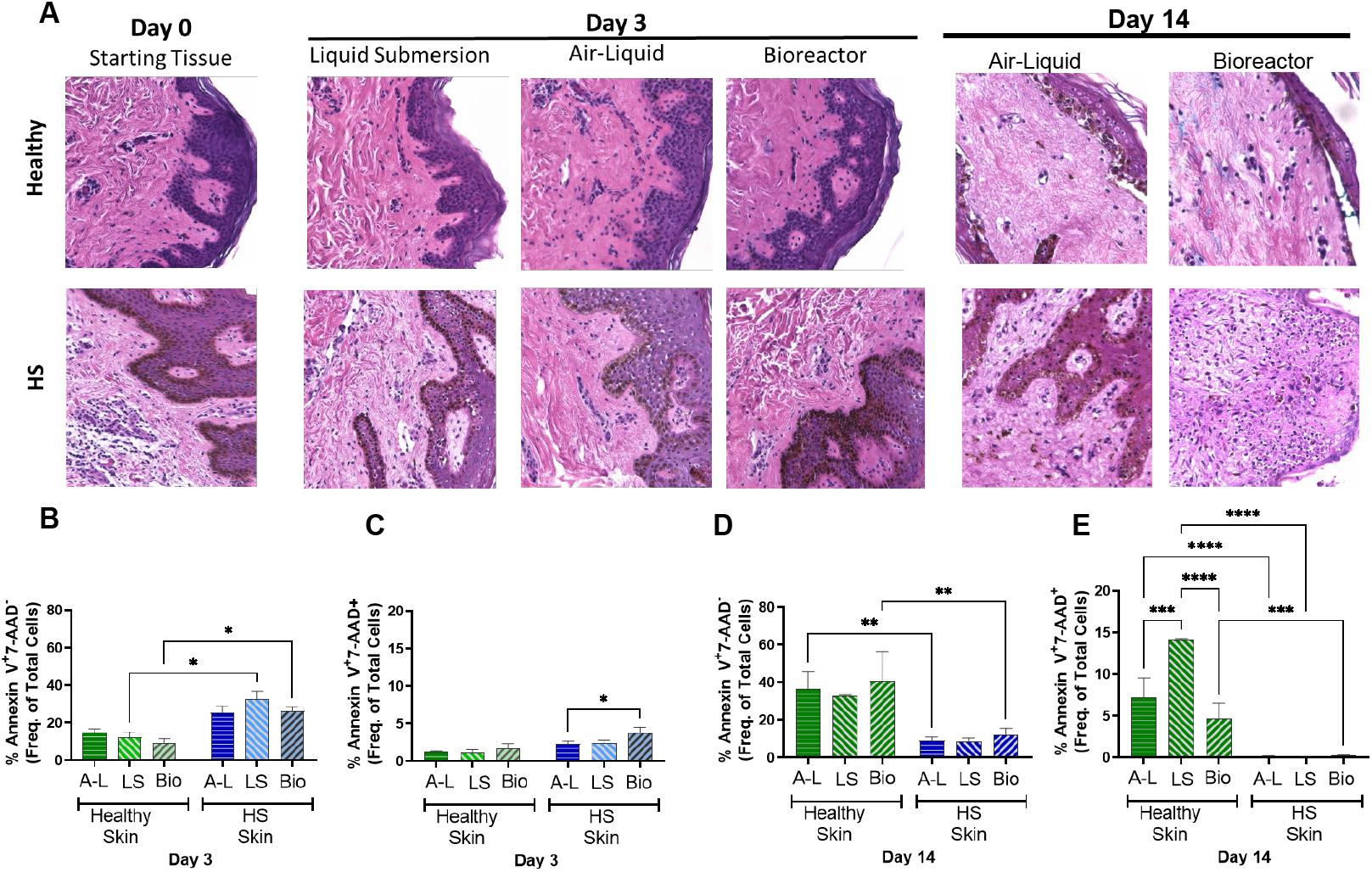
Tissue Histology to Demonstrate Architecture and Viability following *Ex Vivo* Culture. **A.** Histologic architecture of *ex vivo* cultured tissue is similar to that observed in starting tissue for both healthy and HS skin. **B-C.** When early apoptosis (**B)** and late apoptosis/necrosis (**C**) were evaluated via flow cytometry, HS skin was found to have a higher proportion of early apoptotic cells when compared to healthy skin at day-3 with L-S and Bio culture. The proportion of late apoptotic/necrotic cells did not differ between healthy and HS skin, although an increase was seen with Bio culture of HS skin **D-E**. Following 14 days culture, early apoptosis (**D**) and late apoptosis/necrosis (**E**) were increased in healthy skin. Additionally, L-S culture of healthy skin increased late apoptotic/necrotic cells when compared to the other two culture platforms. n=3-14. *p≤0.05, ** p≤0.01, *** p≤0.005, **** p≤0.0001.

### Apoptosis analysis

Flow cytometry data did not show significant differences in early apoptosis among the three culture platforms up to day-3. However, the percentage of early apoptotic cells was higher in HS skin culture (L-S: 32.8 ± 4.26, Bio: 26.4 ± 2.14) relative to healthy skin (L-S: 12.2 ± 2.9, Bio: 9.47 ± 2.1; p=0.019 L-S culture, p=0.017 Bio culture) (**Figure 2B**). No significant differences were observed in the percentage of late apoptotic/necrotic cells among the three platforms of healthy skin cultures, but late apoptotic/necrotic cells were significantly higher in Bio culture (3.83 ± 0.69) of HS skin relative to A-L culture (2.29 ± 0.4; p=0.026) at 3 days (**Figure 2C**). Interestingly at day-14, a moderate increase in apoptotic cells was observed the healthy skin cultures relative to HS skin cultures. When healthy and HS skin cultures were compared, a significant increase in apoptotic cells was observed in A-L culture (36.37 ± 9.27) and bioreactor culture (40.8 ± 15.7) from healthy skin when compared to A-L culture (8.9 ± 2.26) and bioreactor culture (12.18 ± 3.2) of HS skin (A-L culture: p=0.0074, Bio culture: p=0.0069) (**Figure 2D**). Similarly, an increase in the late apoptotic population stained with Annexin V^+^ 7-amino-actinomycin D^+^ (7-AAD) was observed at day-14 relative to day-3 of healthy skin culturs. L-S culture of healthy skin contained the highest proportion of late apoptotic cells (14.5 ± 1.83) when compared to A-L culture (7.17 ± 2.38; p=0.0001) and Bio culture (4.68 ± 0.15; p<0.0001). Additionally, when healthy and HS skin culture were compared, a significant increase in the late apoptotic population was observed in healthy skin samples when using all culture platforms (**Figure 2E**).

### Characterization of various cell populations

T cell populations were evaluated following 3 days of culture. Although the CD4^+^ population was very high in HS skin (11.27 ± 3.86) relative to healthy skin (2.95 ± 0.168; p=0.037) at day-0, no significant difference was observed in proportion of CD8^+^ T cells between healthy and HS skin (**Figures 3A and 3B**). Upon further analysis, bioreactor culture was shown to best represent the CD4^+^ T cell populations observed in the starting tissue of healthy skin, while a significant decrease in this population found with A-L (2.92 ± 1.08; p=0.011) and L-S (2.65 ± 0.93; p=0.011) culture at day-3 relative to HS skin at day-0 (11.27 ± 3.86). Interestingly, CD4^+^ tissue resident memory (TRM) cells, regulatory T cells (T_reg_), and central memory T cells were expanded in all culture platforms in HS specimen when compared to their levels in the starting tissue (**Figure 3C**). Whereas TRM, effector and central memory T cells were decreased in healthy skin. When CD8^+^ T cell populations were compared in healthy skin, the A-L and L-S culture platforms were found to best maintain similar proportions to those seen in the starting tissue. However, T_reg_ increased and TRM and central memory T cells decreased when using all culture platforms (**Figure 3D**). In HS skin, CD8^+^ T cell populations remained similar to the starting tissue across culture platforms however slight increases in T_reg_ and central memory T cells, as well as slight decreases in effector memory cells were seen with culture. When activated T cell populations were evaluated, a significant increase in activated CD4^+^ T cells (AL culture: 16.83 ± 4.37, p=0.0001; L-S culture: 12.03 ± 2.07, p=0.035; Bio culture: 12.07 ± 3.19, p=0.034) was observed following culture of HS skin tissues at day-3 relative to healthy skin tissues at day-0 (2.94 ± 0.62) (**Figure 3E**). Similarly, activated CD8^+^ T cells were elevated with culture but the number did not reach significance in either tissue type or across culture platforms (**Figure 3F**). Proliferating CD4^+^ T cells were increased with Bio culture in HS tissue, but no other changes were observed and no changes in proliferating CD8^+^ T cells were found (**Figures 3G and 3H**).

**Figure 3:**
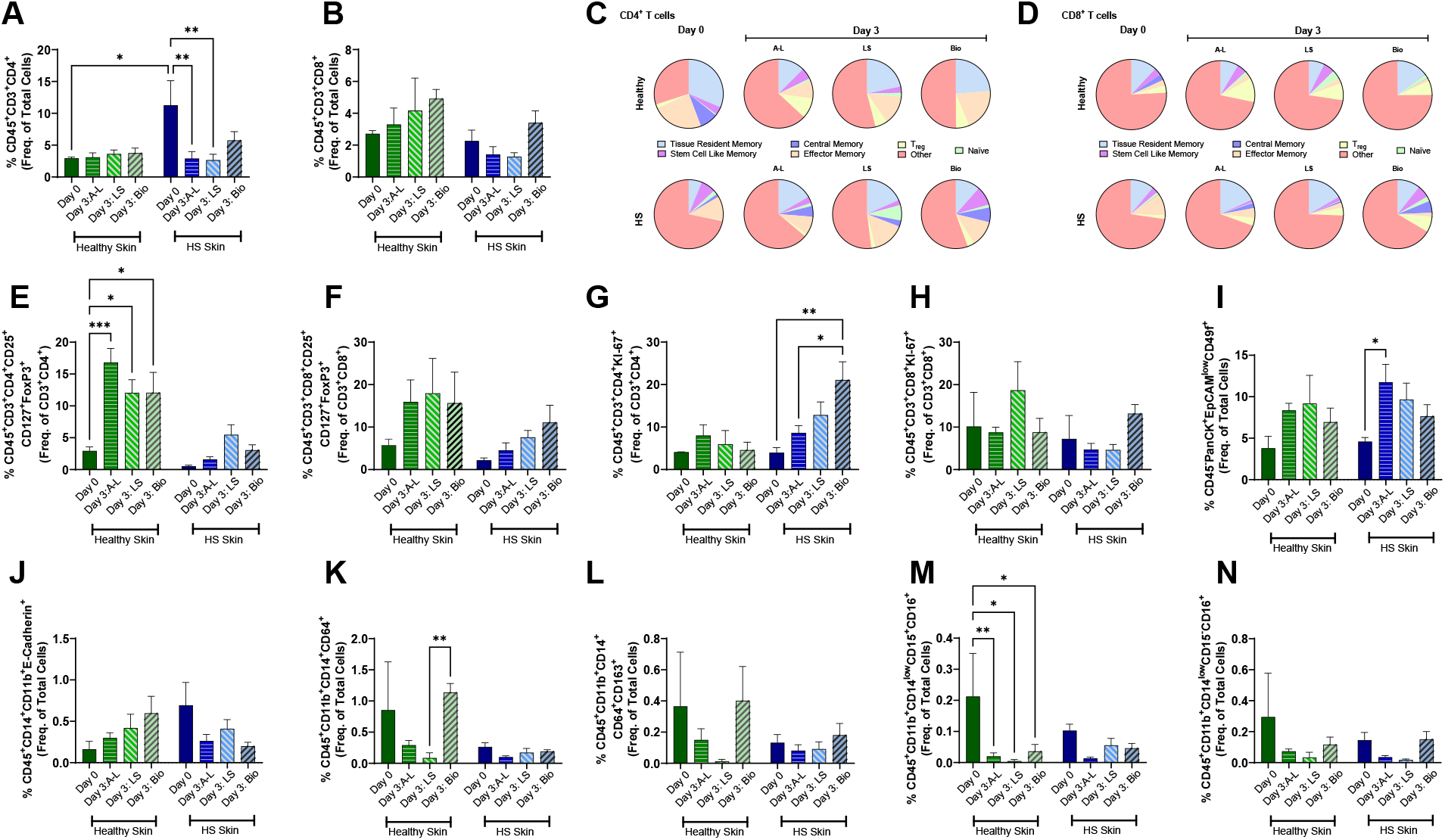
Cellular Heterogeneity is Maintained with *Ex Vivo* Culture. **A-B.** CD4^+^ T cells are increased in HS skin when compared to healthy skin at day-0, but loss of this population is seen with culture (**A**), whereas CD8^+^ T cells are maintained (**B**). **C-D.** Subpopulations of CD4^+^ (**C**) and CD8^+^ (**D**) T cells vary somewhat with culture platform. **E-F.** Activated CD4^+^ T cells increased with culture in healthy skin (**E**), whereas no significant change was observed in activated CD8^+^ T cells (**F**). **G-H.** Proliferating CD4^+^ T cells (**G**) were increased with Bio culture in HS skin, whereas no changes were observed in CD8^+^ T cells (**H**). **I.** The proportion of keratinocytes was similar between healthy and HS skin, however an increase in this population was observed with A-L culture of HS skin. **J**. Langerhans cells showed no significant change between healthy and HS skin or with culture. **K-L.** The proportion of macrophages (**K**), M2 like macrophages (**L**), neutrophils (**M**), and natural killer (NK) cells (**N**) were small in both healthy and HS skin. n=3-12. *p≤0.05, ** p≤0.01.

When the proportion of keratinocytes and proliferating keratinocytes were evaluated, a significant increase in keratinocytes was found in A-L culture (11.74 ± 2.17) relative to the starting tissue (4.6 ± 0.48) in HS skin (p=0.035), however no other significant changes were observed between tissue type or across culture platforms (**Figures 3I and Sup. Figure 1A**). Similarly, no changes were observed when the proportion of Langerhans cells was assessed (**Figures 3J and Sup. Fig 1B**). Although the proportion of macrophages and proliferating macrophages were small within both healthy and HS skin specimen, these cell populations were found to increase with bio culture in healthy skin (**Figures 3K and Sup. Fig 1C**). When M2 macrophages and proliferating M2 macrophages were assessed no significant changes were observed (**Figures 3L and Sup. Fig 1D**). Similarly, the proportions of neutrophils (**Figure 3M**) and natural killer (NK) cells (**Figure 3N**) were found to be small within both healthy and HS skin. While no significant differences were observed in the NK cell population, a significant decrease in neutrophils was observed with all culture platforms in healthy skin tissue when compared to the starting tissue.

### Status of inflammatory markers genes in *ex vivo* HS models

Next, we carried out mRNA expression profiling of 586 inflammatory genes using TaqMan based human inflammation open array panel to assess the dynamic regulation of the expression of inflammatory genes in these culture models. Data collected from i0nflammatory arrays demonstrate that HS skin tissue maintain the dynamics of the inflammatory signatures in *ex vivo* skin culture platforms (**Figure 4A**), although, these changes were not identical across three culture platforms. The expression of many inflammatory signatures were progressively elevated at day-14, as shown by the heat map (**Figure 4A**). Pathway analysis using Ingenuity Pathway Analysis (IPA) showed that the osteoarthritis pathway, IL6 signaling, the pathways associated with the role of IL-17A in psoriasis, acute phase response signaling, STAT3, NFκB and TLR signaling remained elevated in all *ex vivo* culture platforms up to day-14 (**Figure 4B**). On the other hand, downregulated gene signatures affected pathways such as LXR/RXR activation, PPARα/RXRα activation, antioxidant action of vitamin C, CREB Signaling in Neurons, PI3K/AKT signaling in the pathogenesis of influenza, PTEN signaling, phagosome formation, Eicosanoids signaling and crosstalk between dendritic and NK cells (**Figure 4C**). Many pathways were unchanged in culture at all time points, including G1/S checkpoint regulation pathway, Gαi signaling, apoptosis signaling, coagulation system, endocannabinoid cancer inhibition pathway and the ferroptosis signaling pathway (**Figure 4D**).

**Figure 4:**
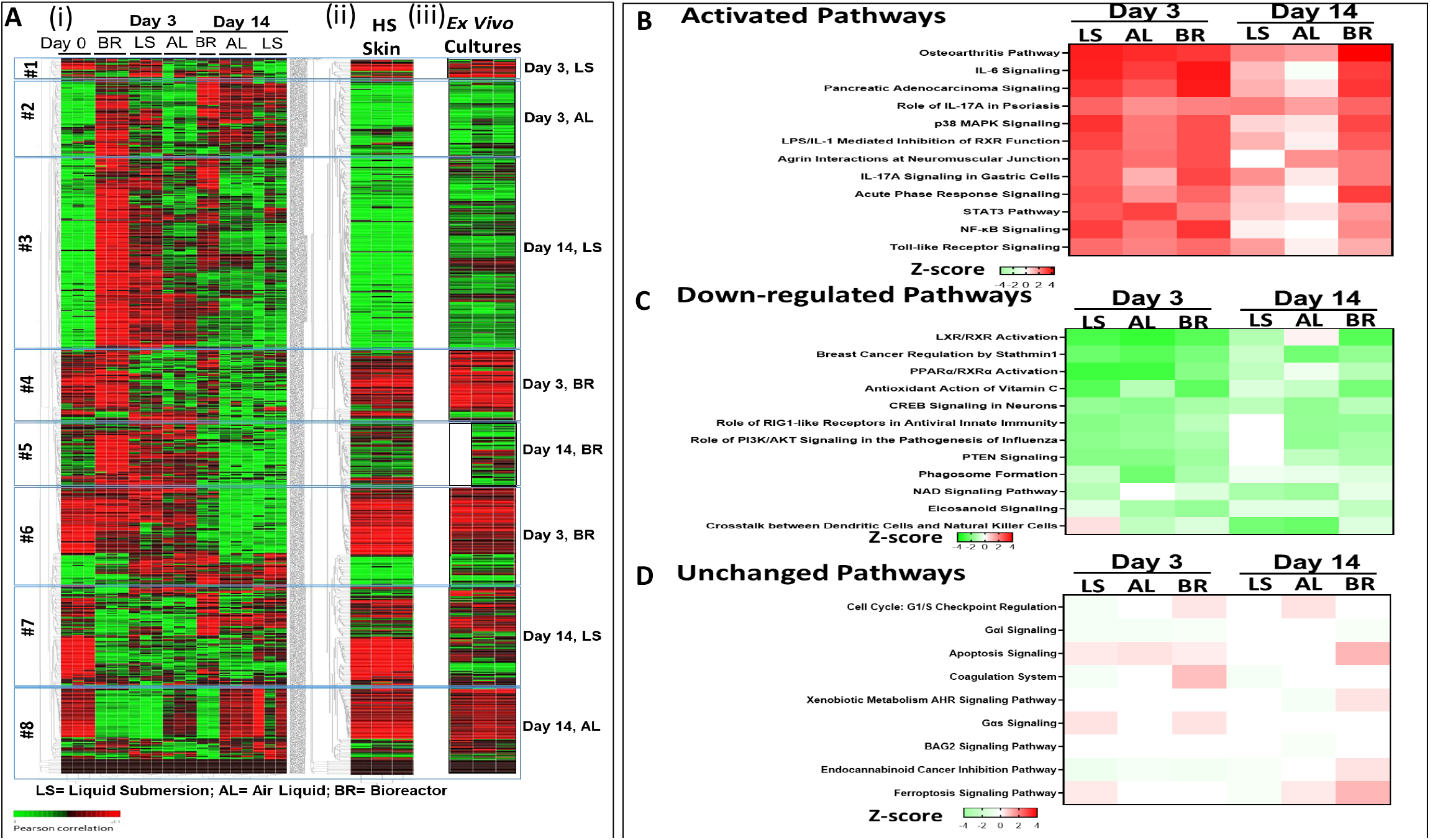
Comparison of inflammatory genes profile of HS skin before and after organ culture for upto fourteen days. Based on gene expression profile, we divided heatmap into eight gene clusters. This demonstrate that any of the three different culture methods alone is not sufficient but these three together provide great value as a tool to understand molecular pathogenesis of HS and its treatment, we stitched clusters that match with HS skin at day-0. As can be seen that four of the day-3 clusters and four of the day-14 clusters from three different culture method match. Thus, a fourteen-day drug treatment to demonstrate its efficacy is possible under these conditions. **A (i)** Heatmap showing the change in expression of various inflammatory genes in three different *ex vivo* culture platforms at day-3 and day-14 of HS skin culture relative to inflammatory signature at day-0 of HS skin excision. **(ii)** Heat Map of HS skin at day-0. **(iii)** Stitched heat Map derived from the most resemble segment of heatmaps of various *ex vivo* cultures either at day-3 or day-14 after dividing all the heat maps into 8 segments. **B.** Heat Map showing upregulated pathways at all time points of *ex vivo* culture of HS skin, relative to HS skin at day-0. **C.** Heat Map showing downregulated pathways at all time points of *ex vivo* culture of HS skin, relative to HS skin at day-0. **D**. Heat Map showing pathways remained unchanged during *ex vivo* culture of HS skin, relative to HS skin at day-0.

The inflammatory pathways, such as hepatic fibrosis signaling, cardiac hypertrophy signaling, production of NO and ROS in macrophages, dendritic cell maturation, TEC kinase signaling, HER-2 signaling in breast cancer, ILK signaling, Type I Diabetes mellitus signaling, iNOS signaling, HMGB1 signaling and IL9 signaling **Figure 5A**), were upregulated at day-3 in various cultures but downregulated and/or returned to basal levels by day-14 of culture. The signaling pathways, which remained highly activated at day-14 of various *ex vivo* HS skin cultures were the osteoarthritis pathway, the pathway associated with the role of IL-17A in psoriasis, and LPS/IL-1 mediated inhibition of RXR function. Interestingly, Fcγ receptor-mediated phagocytosis in macrophage and monocytes was only upregulated in various *ex vivo* culture at day-14 (**Figure 5B**). Overall, the data from the inflammatory array demonstrate that these *ex vivo* HS skin cultures are suitable to defining the molecular pathogenesis of inflammation in HS skin.

**Figure 5:**
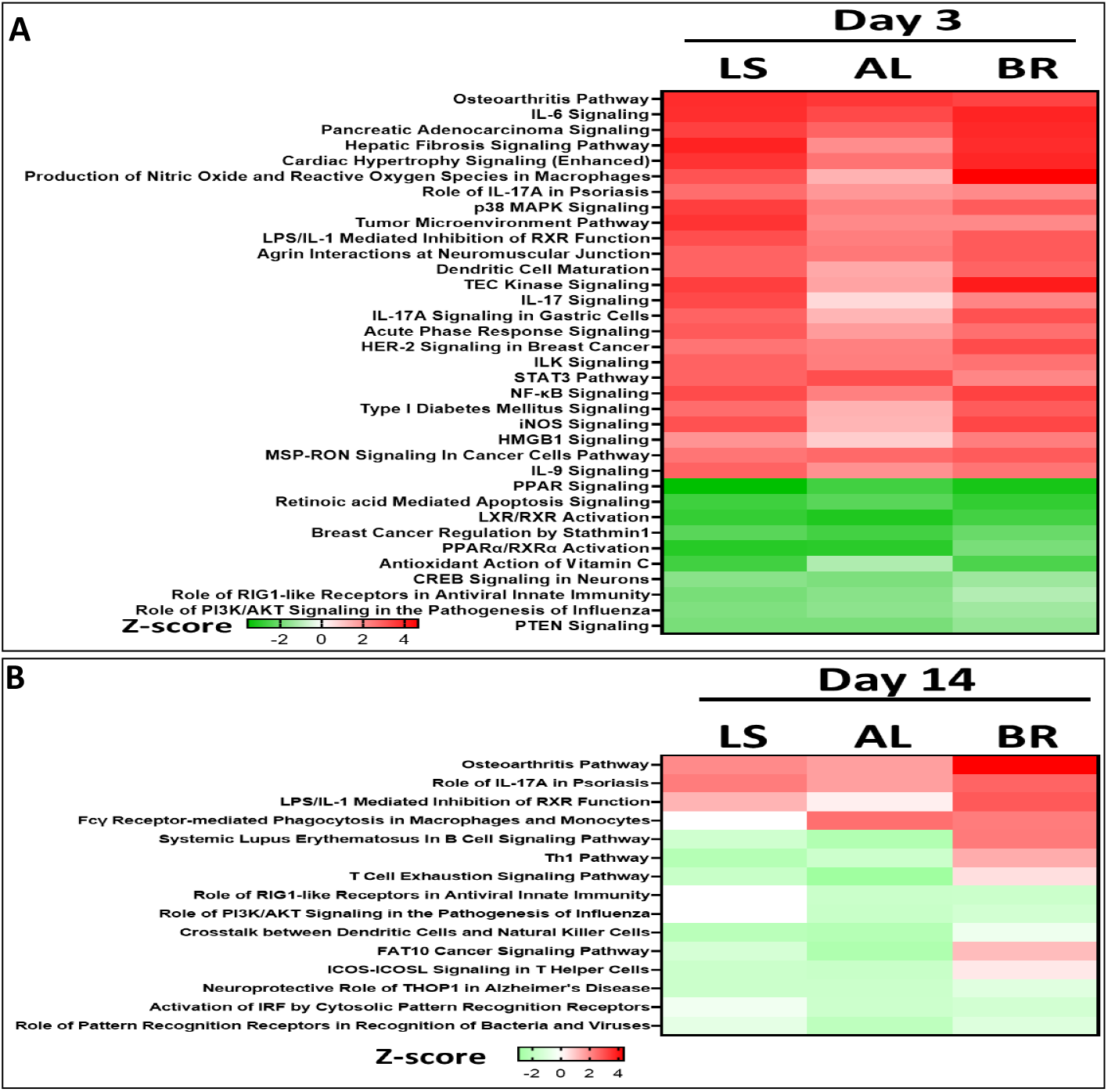
IPA Analysis Showing that Most Inflammatory Pathways were Upregulated at Day-3 of *Ex Vivo* Culture, but Few could be Sustained at Day-14. **A.** Heat Map showing the pathways significantly upregulated or downregulated at day-3 of *ex vivo* culture. **B**. Heat Map showing the pathways significantly upregulated or downregulated at day-14 of *ex vivo* culture.

### Verification of suitability of *ex vivo* HS skin culture models for drug screening

To verify the suitability of these *ex vivo* skin cultures to predict treatment efficacy, we employed an immunomodulatory drug, lenalidomide, and a bromodomain and extra-terminal (BET) inhibitor, CPI-0610, in all three *ex vivo* culture platforms for 3 days of culture. Both of these drugs are expected to alter inflammatory profile by different mechanisms ^18,19^. The histopathologic evaluation using H & E staining did not reflect major differences in tissue architecture due to drug treatment and associated toxicity in healthy skin when using any of the three culture methods (**Figure 6A & Sup. Figure 2**). Moreover, no significant change in early apoptotic (Annexin V^+^ 7AAD^-^) or late apoptotic/necrotic (Annexin V^+^ 7AAD^+^) cell populations was observed between drug treated - and vehicle-treated groups using any of all these culture platforms at day-3 time point (**Figure 6B**). However, an overall increase in late apoptosis/necrosis in HS tissues was observed. None-the-less, we did not established a dose response relationship for any of these drugs at this stage (**Figure 6C**).

**Figure 6:**
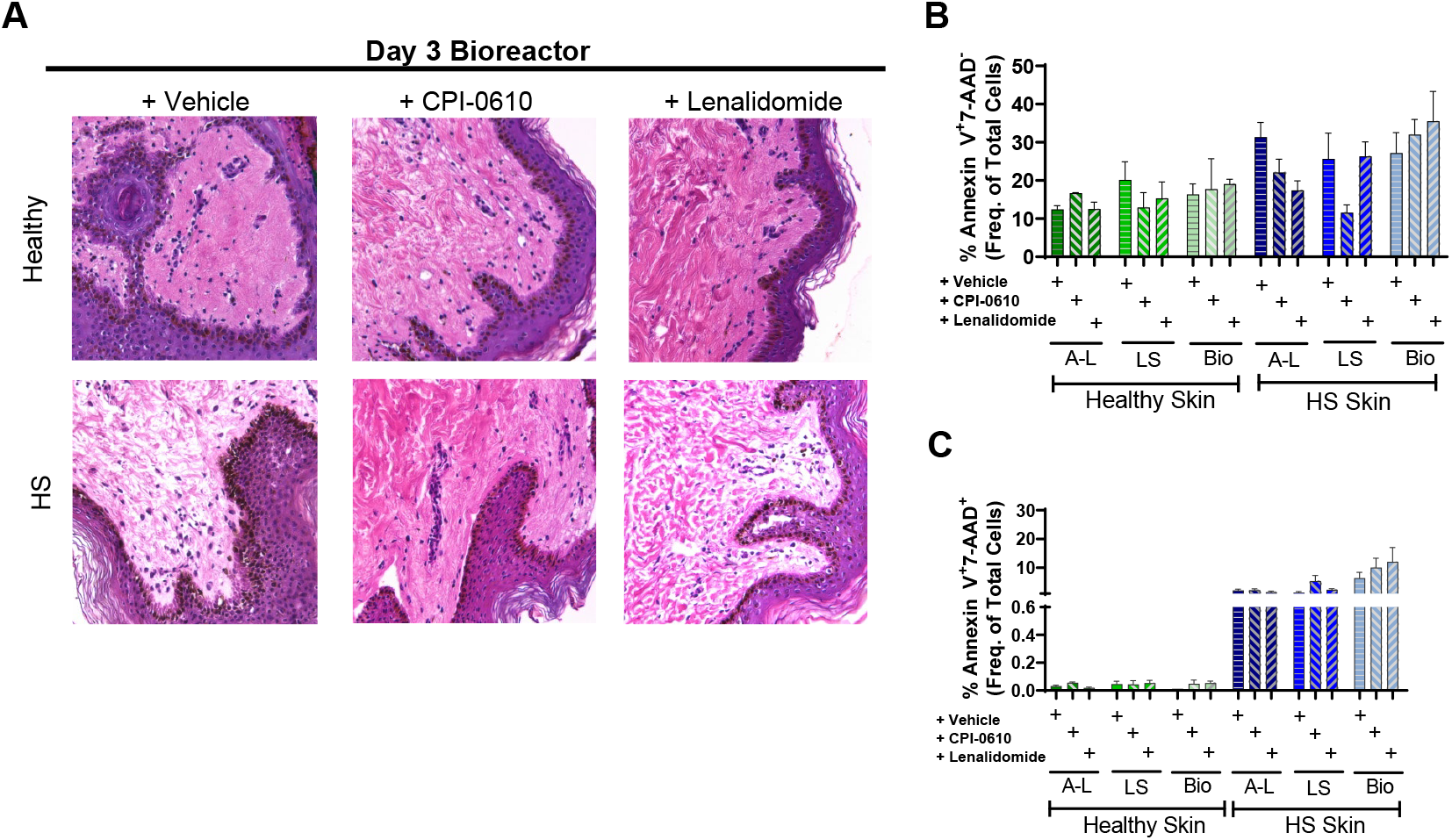
Effects of Drug Treatment on Tissue Histologic Architecture and Viability following *Ex Vivo* Culture. **A.** Histologic architecture of *ex vivo* cultured tissues treated with lenalidomide or CPI-0610 is similar to that observed in vehicle control treated tissue for both healthy and HS skin when cultured using the Bio platform. **B.** When early apoptosis was evaluated via flow cytometry, no significant difference was noted with treatment or culture platform in healthy or HS skin. **C** Late apoptosis/necrosis was increased in HS skin when compared to healthy skin, however no changes in response to treatment or culture platform were observed. n=4 per group.

Employing Luminex analysis for various cytokines/chemokines and growth factors in the supernatant procured from drug- and vehicle-treated cultures, we observed that both drugs effectively blocked the secretion of various cytokines/chemokines and growth factors in the supernatants (**Figure 7**). Both drugs were able to reduce the levels of Th1/Th2/Th9/T_reg_ associated cytokines such as IL2, IL4, IL5, IL9, IL21; while inflammatory cytokines/chemokines such as IL8, IFNα, MIP1α, SDF1α, MCP1 GROα/KC *etc*. and various growth factors namely, PDGF-1, VEGFd, and SCF, were also diminished in the supernatant procured from air-liquid *ex vivo* HS skin cultures at day-3 relative to vehicle-treated controls (**Figure 7A**). Secretion of some of inflammatory cytokines/chemokine viz., MIP1α, IL-1RA, IL6 and IFNα were also blocked in the supernatant of ex vivo HS skin from bioreactor culture at day-3 of culture (**Figure 7B**). However, only CPI-0610 was effective in blocking many cytokines/chemokines and growth factors, as assessed in the supernatant of *ex vivo* HS skin tissue of liquid-liquid culture at day-3 with exception to IL2, IL4, MCP1, IL9 and VEGFd (**Figure 7C**). Interestingly, these were not inhibited in the supernatant of air-liquid *ex vivo* HS skin cultures at day-3. Multiple studies have also evaluated cytokines and other inflammatory proteins in lesional, peri-lesional skin wound exudate, and serum of HS patients to find out the importance of these proteins in the pathogenesis and treatments ^20–23^.

**Figure 7:**
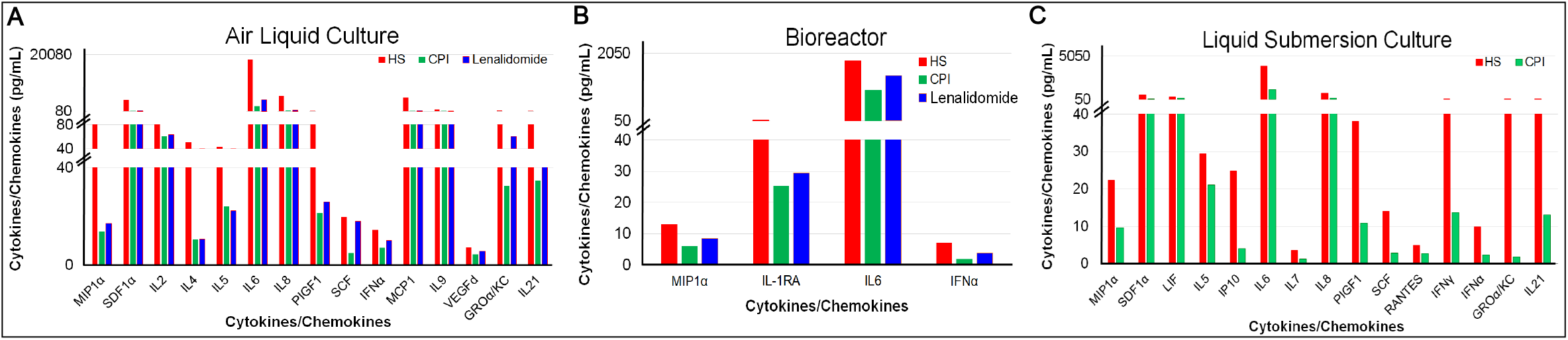
Effect of drug treatment on anti-inflammatory cytokine secretion in culture supernatant of E*x vivo* Models of HS. Changes in the levels of various cytokine/chemokines and growth factors in supernatant procured from either Vehicle (HS), CPI-0610 (CPI), or lenalidomide treated *ex vivo* HS skin models from Air-Liquid Interface (**A**), Bioreactor (**B**) and Liquid submersion (**C**) platforms.

Reconstructing a heat map from these model cultures was achieved by dividing the heatmap of HS day-0 skin detailing the expression profile of inflammatory genes in eight clusters, as shown in (**Figure 4A** (iii)). Thereafter, we searched the expression profiles of all three cultures at day-3 and day-14, and identified the segments where we did not find any alteration in the expression of the clusters compared to day-0 profile. These unaltered segments were sutured to make an artificial heatmap, as shown in **Figure 4A** (iii). This heatmap could be utilized to predict the impact of drug treatment on the profile of inflammatory genes in HS skin (day-0). In sum, this reconstruction of the heatmap was necessary for drug evaluation, as our culture methods provide a dynamic environment for the gene expression and many of these genes are continuously altered.

Overall, various *ex vivo* cultures together provide a powerful system to assess the drug effects as exemplified by demonstrating the effect of two different class of mechanistically anti-inflammatory drugs.

## Discussion

In this study, we established three different *ex vivo* culture model systems for healthy and HS skin and assessed suitability of these *ex vivo* models to screen drugs which can affect inflammatory signaling and progression of inflammation in HS skin. Previously, *Vossen and colleagues used* Transwell *ex vivo* skin culture from healthy and HS skin ^24^. Using these cultures authors showed that this methodology could be used for *ex vivo* evaluation drug screening of HS tissue. In this Transwell culture, following exposure HS skin tissue to prednisolone or biologics targeting TNF-α, interleukin (IL)-17A, IL-12/23p40 or CD20 (adalimumab, infliximab, secukinumab, ustekinumab and rituximab, respectively), TNF-α inhibitors and prednisolone were found to be the most powerful inhibitors of proinflammatory cytokines. In comparison to the previous attempts to establish skin tissue cultures, our three different types of culture platforms namely, air-liquid interface, liquid submersion and a more controlled perfusion bioreactor, establish *ex vivo* culture of healthy and HS skin and provide a more powerful and rigorous tool in this regard. In an attempt to demonstrate the suitability of these cultures for understanding the molecular pathogenesis of disease progression, particularly with respect to the dynamic regulation of inflammatory cascade, we characterized various cell populations including keratinocytes and immune cells in these cultures. Our initial findings demonstrated that these *ex vivo* cultures were able to maintain a significant level of the tissue architecture and integrity at least up to day-3 of culture. Furthermore, when cultures were extended to day-14, the tissue architecture and integrity were maintained in HS skin culture but not in healthy skin culture. At this stage to maintain identical culture conditions and to avoid any complexity in explaining the need for different culture conditions, we did not try to alter the culture conditions of healthy skin to maintain it for longer duration of time.

These results are primarily based on H & E staining of the cultured skin. However, a further detailed analysis is needed to assess each cell-type and their ability to maintain the molecular signatures, for example, using single cell analyses. The observed differences in healthy and HS skin cells may be due multiple factors, particularly due to differences in metabolic signatures. In this regard, Vitamin D metabolism, steroid metabolism, biological processes associated with collagen and extracellular matrix remodeling, growth and tissue factors such as tissue factor 3 (T3), growth and differentiation factor 2 (GDF2), midkine and clusterin which are known to play important roles in vital cellular and physiological processes and are dysregulated in HS skin ^25–27^. The compromised metabolic environment of HS skin is considered to be associated with HS disease progression ^4,28^

Next, evaluation of various cell populations in these *ex vivo* cultures showed that bioreactor culture at day-3 was able to best represent the CD4^+^ T cell populations seen in the starting HS tissue at (day-0). However with further evaluation of CD4^+^ T cells, TRM, SCM and central memory T cells were found to expand in all culture platforms in HS specimen when compared to the starting (day-0) tissue. CD8^+^ T cell populations remained unaltered when compared to the starting tissue across all culture platforms; however, small increases in T_reg_ and central memory T cells, as well as decreases in effector memory cells was seen. When activated T cell populations were evaluated, an increase in activated CD4^+^ T and CD8^+^ T cells was observed in all culture platforms, but significance could only be achieved for CD4^+^ T cells in *ex vivo* cultures of healthy. Moreover, proliferating CD4^+^ T cells were increased with all the platforms, but significance could be achieved with Bio culture in HS tissue. The ratio of proliferating CD8+ T cells were higher in bioreactor culture of HS skin **(Figure 3)**. Evaluation of keratinocytes and proliferating keratinocytes showed that these cultures only support the healthy growth of epidermal cells upto day-3 of evaluation but hyper-proliferation of keratinocytes was maintained in these *ex vivo* cultures of HS skin day-14. The proportion of macrophages and proliferating macrophages were small within both healthy and HS skin specimen, and these cell populations were found to increase with bio culture in healthy skin. However, the proliferating M2 population was found to be decreased in A-L and L-S culture platforms of HS skin. Overall, these data demonstrate that these *ex vivo* models were not only able to maintain various cells populations but were also able to preserve the disease tissue system physiology at least partially. The further details and other mechanistic aspects remain to be investigated and are beyond the scope of this study.

Molecular profiling of inflammatory genes in these culture showed that there are dynamic expression profiles and a single culture platform cannot represent the inflammatory status of HS skin at day-0. However, using all three *ex vivo* culture platforms at day-3 and day-14 could recapitulate most of the inflammatory pattern seen in HS skin. Molecular expression data from these cultures also indicates that these cultures could serve as useful models to screen anti-inflammatory drugs. In this regard, our data employing the immunomodulatory drug, lenalidomide and the anti-inflammatory/epigenetic modulatory drug, CPI-0610 (a bromodomain and extra-terminal (BET) inhibitor) in these *ex vivo* HS models showed that many inflammatory cytokines/chemokines were downregulated by these treatments. Although, as described earlier, a single culture method was not as powerful as the results obtained from the three culture methods together.

Thus, our combined *ex vivo* culture models provide a unique opportunity to employ these systems to understand the pathobiology of HS as well as screening and development of various therapeutic agents, particularly those which are already FDA approved for other conditions and could be repurposed for HS treatment.

### Conclusion

We believe that these three *ex vivo* culture systems collectively provide a powerful tool for defining the pathobiology of HS and assessing drugs for suppressing inflammatory responses.

## Acknowledgement

This work is supported by NIH grant RO1 ES026219 and NCI grant 5P01CA210946 and by intramural UAB funds to M.A.

## Declaration of no conflict of interest

Authors of this manuscript have no conflict of interest between them or anybody else regarding the scientific contents, financial matters, or otherwise.

---

**Supplemental Figure 1:**
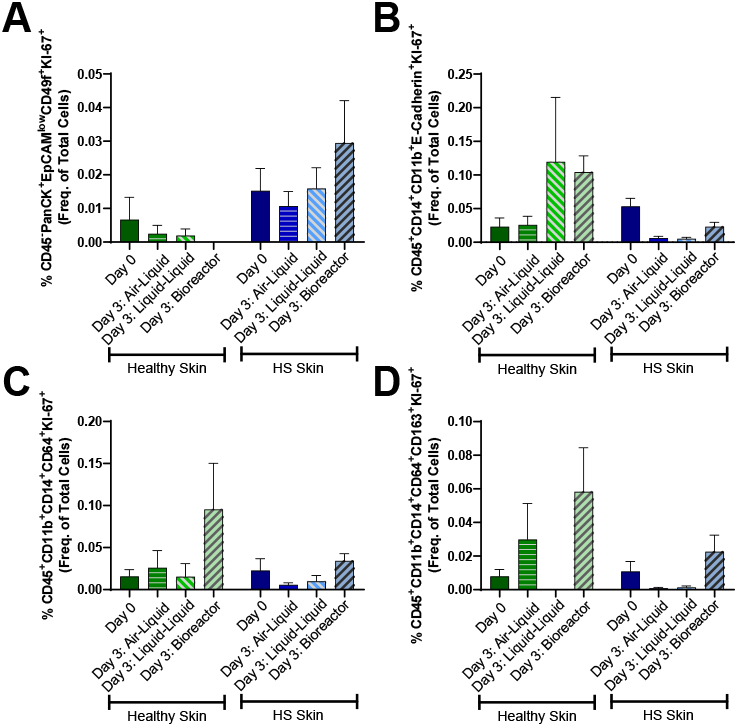
Proliferating Keratinocytes, Langerhans cells, and Macrophage Populations. The proportion of Ki-67^+^ keratinocytes (**A**), Ki-67+ Langerhans cells (**B**), Ki-67+ macrophages (**C**), or Ki-67+ M2 like macrophages (**D**) were not significantly different between healthy and HS skin or across culture platforms. n=3-12.

**Supplemental Figure 2:**
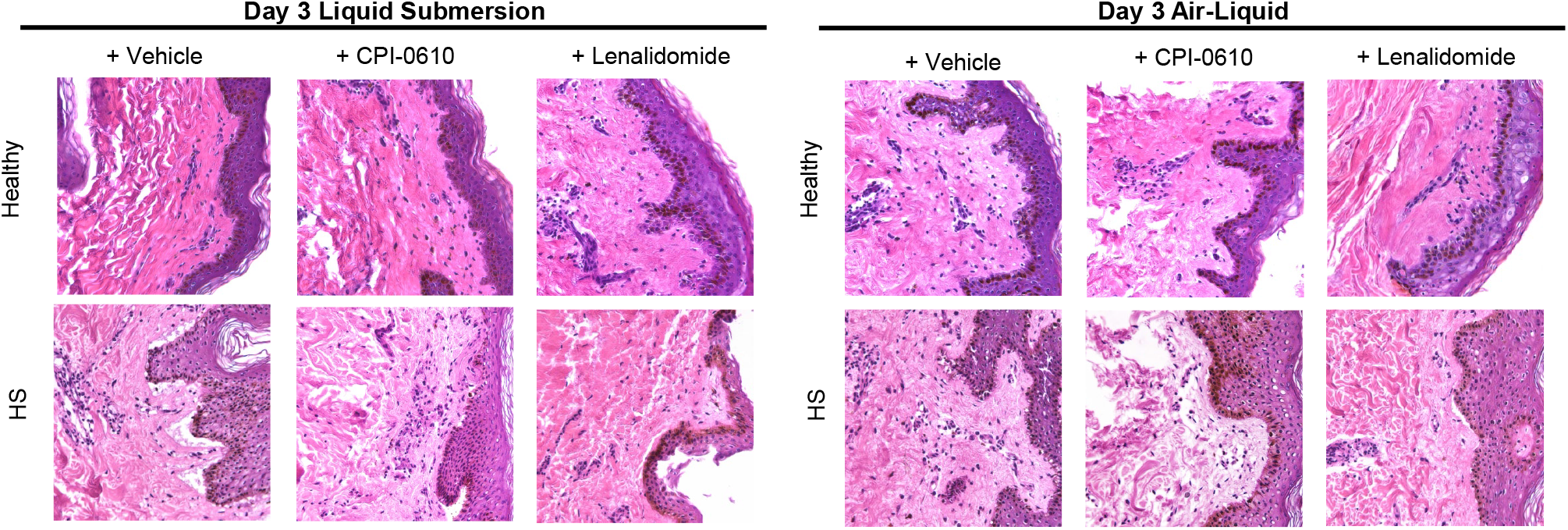
Histologic architecture of *ex vivo* cultured tissues treated with Vehicle, Lenalidomide or CPI-0610 for 3 Days using Liquid Submersion or Air-Liquid Interface Culture. Histologic architecture is similar between CPI-0610, lenalidomide, and vehicle control treated tissues for both healthy and HS skin when using LS (left) and A-L (right) culture.

